# Iron Deficiency Induces Heart Failure with Ectopic Cardiac Calcification in Mice with Metabolic Syndrome

**DOI:** 10.1101/2022.01.20.477010

**Authors:** Yoshiro Naito, Hisashi Sawada, Seiki Yasumura, Tetsuo Horimatsu, Keisuke Okuno, Saki Tahara, Koichi Nishimura, Masanori Asakura, Takeshi Tsujino, Tohru Masuyama, Masaharu Ishihara

**Affiliations:** Department of Cardiovascular and Renal Medicine, Hyogo College of Medicine, Nishinomiya, Japan; Saha Cardiovascular Research Center, College of Medicine, University of Kentucky, KY, USA; Saha Aortic Center, College of Medicine, University of Kentucky, KY, USA; Department of Physiology, College of Medicine, University of Kentucky, KY, USA; Division of Pharmaceutical Therapeutics, Department of Pharmacy, School of Pharmacy, Hyogo University of Health Sciences, Kobe, Japan; Japan Community Health care Organization, Hoshigaoka Medical Center, Hirakata, Japan

**Keywords:** anemia, cardiac calcification, metabolic syndrome, iron deficiency, heart failure

## Abstract

Iron deficiency is linked to worse clinical status and outcomes in heart failure. Although metabolic syndrome contributes to the development of heart failure, the impact of iron deficiency in heart failure complicated with metabolic syndrome remains obscure. KKAy mice were used as a model of metabolic syndrome. Four-week-old male C57BL/6J and KKAy mice were fed either a normal diet or iron-restricted (IR) diet for 12 weeks. During the experiment, 40% of mice died due to pulmonary congestion in KKAy mice with IR diet (KKAy-IR), while no mice died in other groups. Necropsy showed the presence of multiple white lesions on the cardiac surface in those KKAy-IR mice. Echocardiography and histological analyses revealed that KKAy-IR mice exhibited cardiac hypertrophy and cardiac dysfunction with cardiac calcification. Cardiac mRNA of ectonucleotide pyrophosphatase/phosphodiesterase-1 (*Enpp1*), a key enzyme for bone mineralization, was highly abundant in KKAy-IR mice. Of note, iron restriction-induced cardiac calcification and dysfunction were attenuated by etidronate, an inhibitor of bone mineralization, with decreased cardiac *Enpp1* mRNA abundance in KKAy-IR mice. In conclusion, iron deficiency leads to ectopic cardiac calcification and dysfunction with the increase of *Enpp1* in metabolic syndrome model mice.

---

Iron deficiency is associated with poor outcomes in heart failure.^1^ Metabolic syndrome also exerts an injurious role in heart failure as a risk for accelerating its development.^2^ However, the impact of iron deficiency in heart failure complicated with metabolic syndrome remains unknown.

KKAy mice were used as a model of metabolic syndrome as these mice develop obesity, diabetes mellitus, and hyperlipidemia. Four-week-old male wild type (WT) C57BL/6J and KKAy mice were fed either a normal diet or iron-restricted (IR) diet for 12 weeks (n=10/group). While no mice died in other groups, 40% of mice died in KKAy mice with IR diet (KKAy-IR) (**Figure A**). Necropsy revealed the presence of cardiomegaly and pleural effusion in those KKAy-IR mice, indicating cardiac death due to pulmonary congestion (**Figure A**). In addition, multiple white lesions were observed on the cardiac surface. KKAy mice exhibited higher body weight and blood glucose than WT mice, while IR diet did not affect these parameters (**Figure B**). Anemia was observed in mice with IR diet regardless of genotypes (**Figure B**). Serum blood urea nitrogen and creatinine concentrations were comparable among groups (**Figure B**).

**Figure A.**
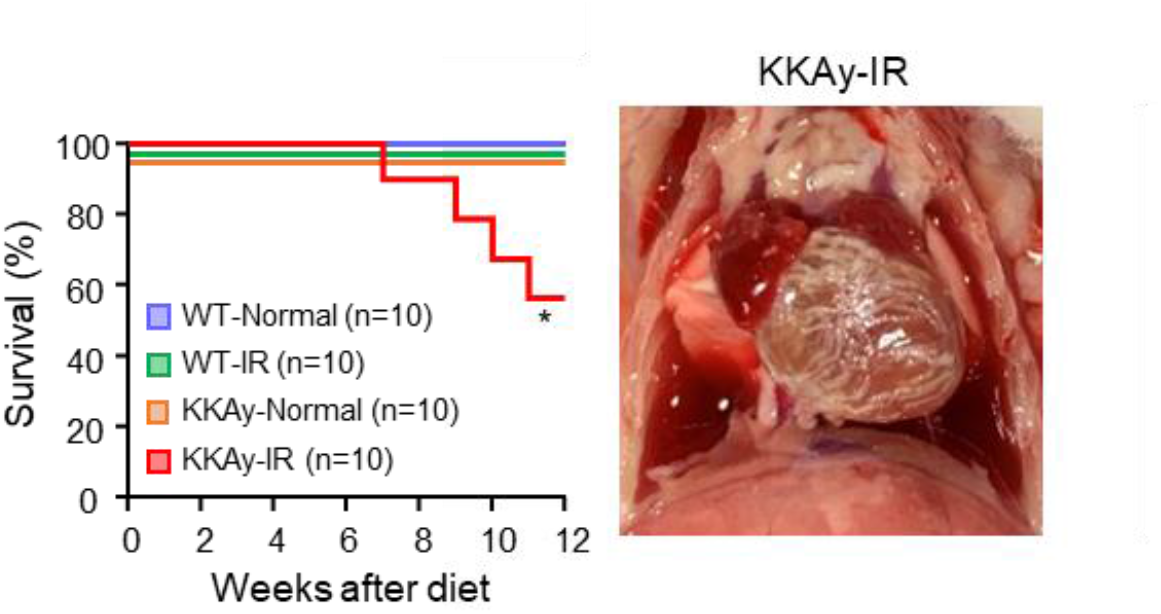
Survival rate in WT mice fed normal diet (blue), WT mice fed iron-restricted (IR) diet (green), KKAy mice fed normal diet (orange), and KKAy mice fed IR diet (red). Representative in situ thorax image of KKAy mice fed IR diet. *P=0.007 vs other groups by Log-Rank test. n=10 per group.

**Figure B.**
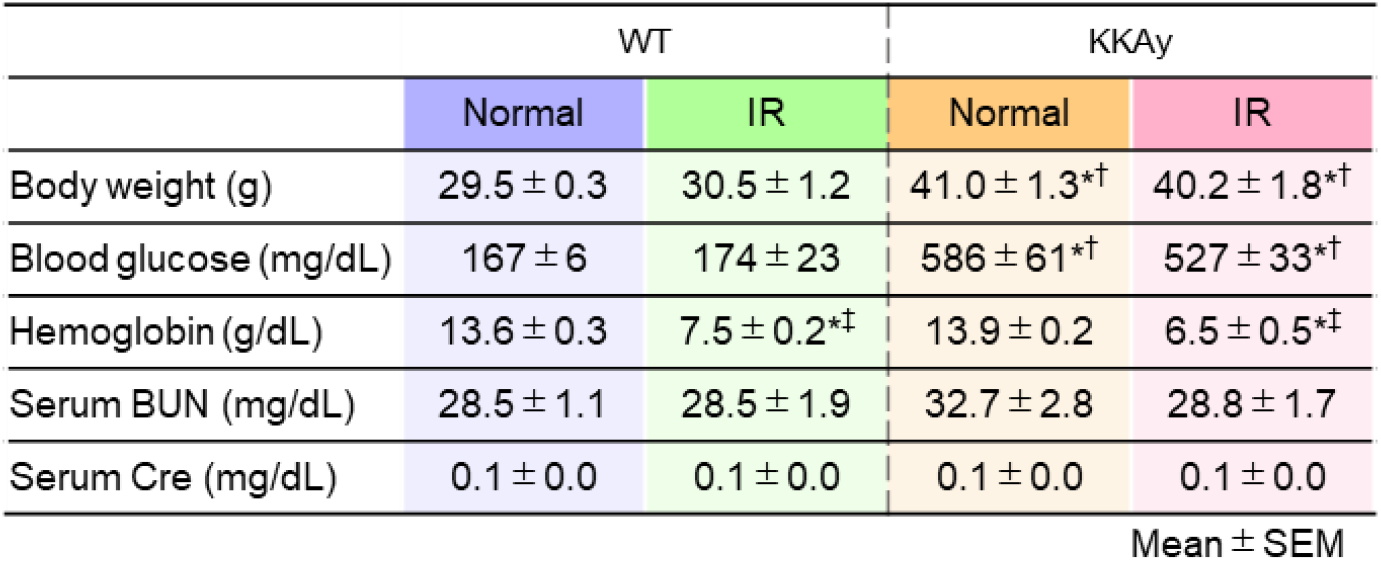
Body weight, blood glucose, hemoglobin, serum blood urea nitrogen (BUN), and creatinine (Cre) concentrations in WT and KKAy mice at 12 weeks after normal or IR diet. n=5-6 per group. *P<0.0001 vs WT mice fed normal diet, ^†^P<0.0001 vs WT mice fed IR diet, and ^‡^P<0.0001 vs KKAy mice fed normal diet.

Ex vivo measurements demonstrated that heart and wet lung weights in KKAy-IR mice were significantly higher than in other groups (**Figure C**). Echocardiography at 11 weeks after diet showed left ventricular dilatation with decreased fractional shortening in KKAy-IR mice (**Figure C**).

**Figure C.**
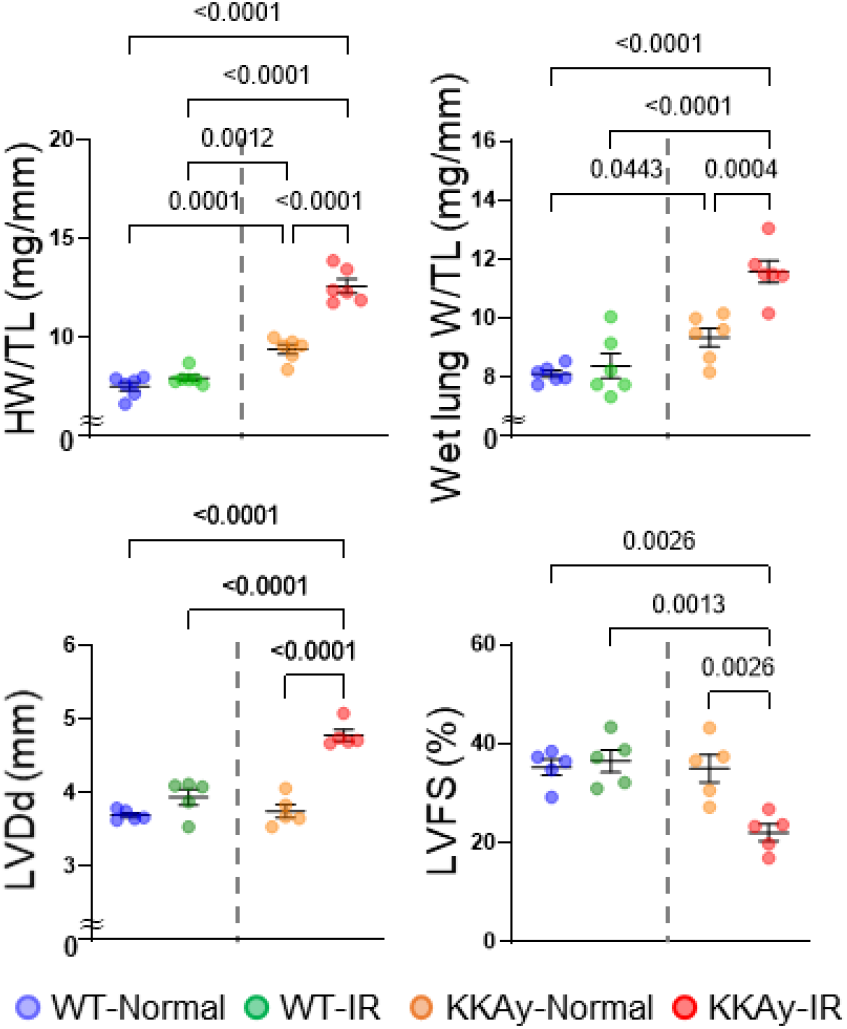
Heart weight/tibia length ratio (HW/TL) and wet lung weight/tibia length ratio (Wet Lung W/TL) in WT and KKAy mice at 12 weeks after normal or IR diet. n=6 per group. Left ventricular end-diastolic dimension (LVDd) and fractional shortening (LVFS) in WT and KKAy mice at 11 weeks after normal or IR diet. n=5 per group.

Ex vivo and histological images revealed that KKAy-IR mice exhibited enlarged hearts and increased cardiac interstitial fibrosis (**Figure D**). Likewise, cardiac mRNA abundance of atrial natriuretic peptide (*Anp*) and collagen type I (*Col1*) were increased in KKAy-IR mice than in other groups (**Figure D**). Subsequently, Alizarin red and von Kossa staining were performed to evaluate cardiac calcification. Multiple calcium depositions were detected in KKAy-IR mice, but not in other groups (**Figure D**). To investigate the molecular mechanism underlying iron restriction-induced cardiac calcification, we assessed osteogenic genes in heart tissues. mRNA of ectonucleotide pyrophosphatase/phosphodiesterase-1 (*Enpp1*), a key enzyme for bone mineralization, was highly abundant in KKAy-IR mice. Runt-related transcription factor 2 (*Runx2*), a key osteogenic transcription factor, was also upregulated in KKAy-IR mice (**Figure D**).

**Figure D.**
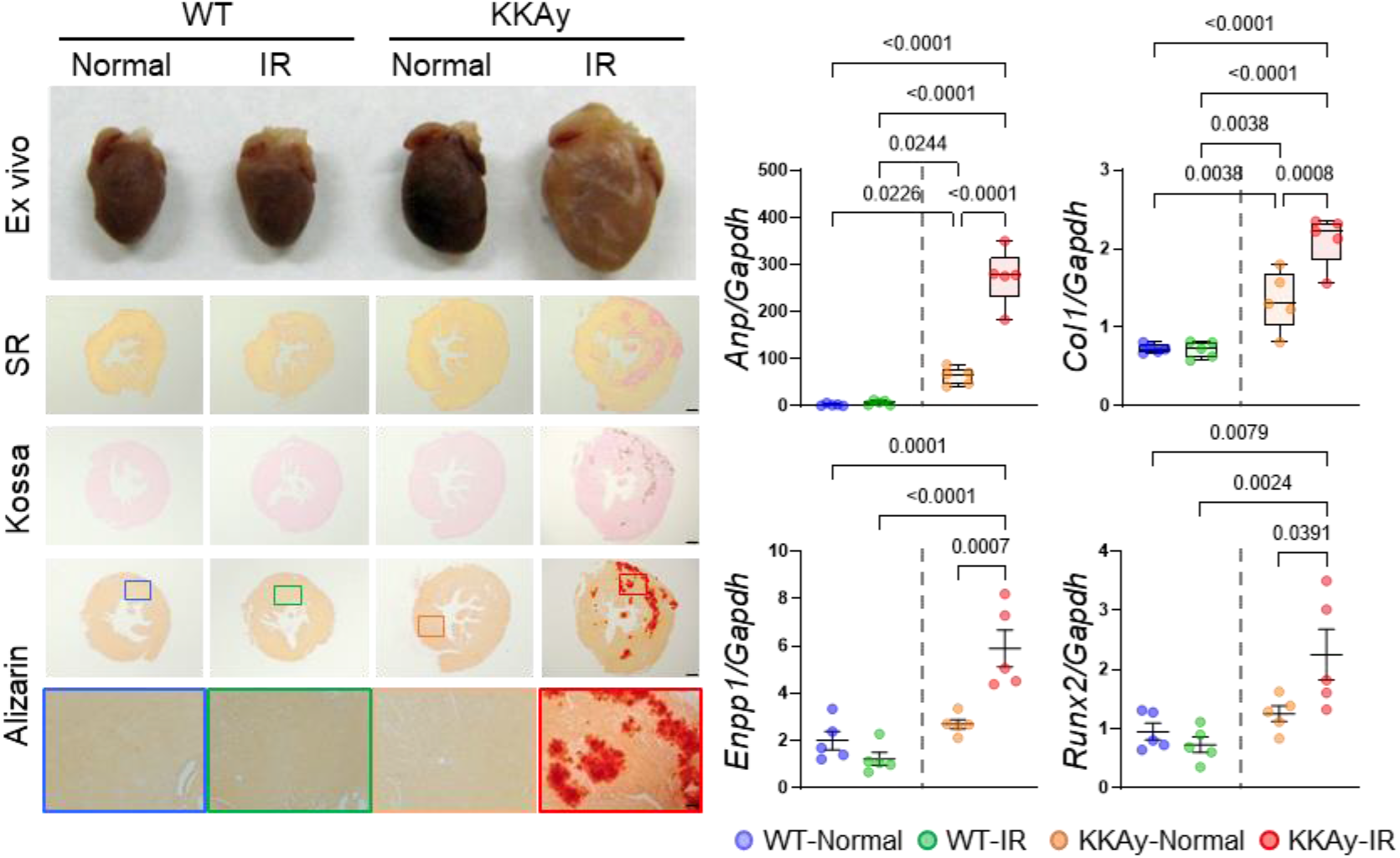
Representative gross appearance of whole hearts and heart sections stained by Picrosirius red (SR), von Kossa, and Alizarin red in WT and KKAy mice at 12 weeks after normal or IR diet (Scale bar: low/high magnification: 500/100 μm). Quantitative analysis of relative *Anp*, *Col1*, *Enpp1*, and *Runx2* mRNA abundance in WT and KKAy mice at 12 weeks after normal or IR diet. n=5 per group.

Next, a time course study was performed to determine the initiation of cardiac calcification in KKAy-IR mice. Heart tissues were harvested from KKAy mice at 3, 7, and 14 days after IR diet. Cardiac calcification was not observed at day 3, while 1 of 6 KKAy mice exhibited mild calcification at day 7. Of note, 5 of 6 KKAy mice (83%) showed mild to moderate calcification at day 14 after IR diet (**Figure E**). Masson’s trichrome staining demonstrated that calcified lesions coincided with collagen deposition (**Figure E**). Cardiac Enpp1 and Runx2 mRNA abundance were increased at day 14 of IR diet compared to 3 days after IR diet (**Figure E**).

**Figure E.**
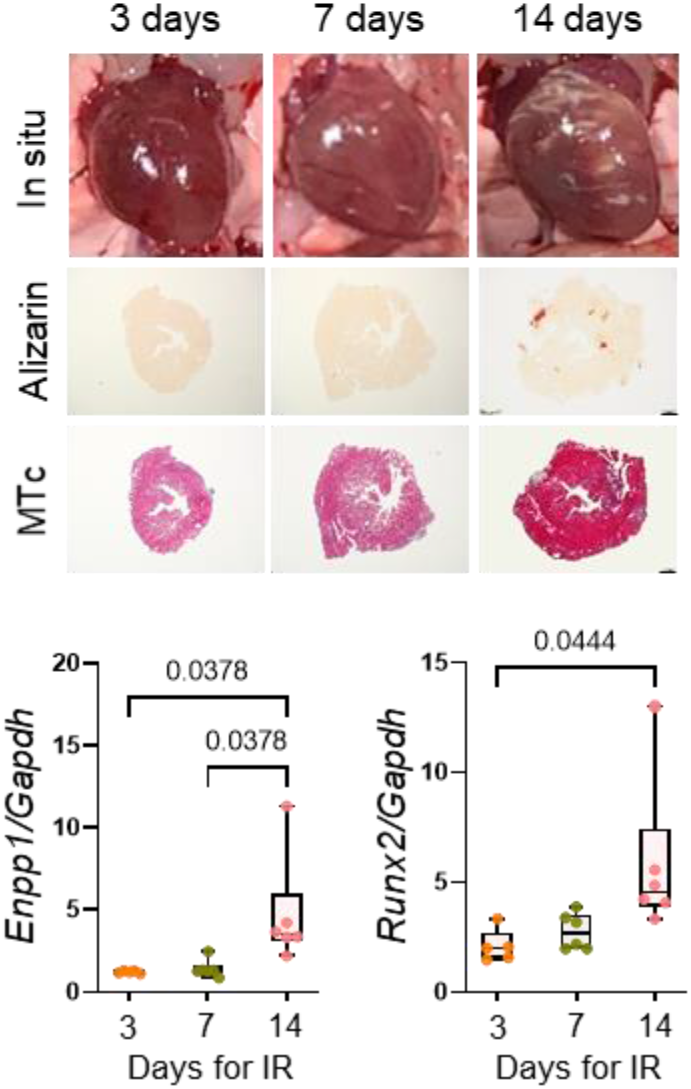
Representative images of in situ heart and heart sections stained by Alizarin red and Masson’s trichrome (MTc) in KKAy mice at 3, 7, and 14 days after IR diet (Scale bar: 500 μm). Quantitative analysis of relative *Enpp1* and *Runx2* mRNA abundance in KKAy mice at 3, 7, and 14 days after IR diet. n=5-6 per group.

Since etidronate, the first bisphosphonate, inhibits bone mineralization, we investigated whether etidronate affects iron restriction-induced cardiac calcification and heart failure in KKAy mice. Either vehicle or etidronate (2.0mg/kg/day, #P5248, Sigma-Aldrich) was administered intraperitoneally for 14 days in WT and KKAy mice. IR diet was started in both groups on the first day of injection. It is noteworthy that cardiac calcification was not detected in etidronate-administered KKAy-IR mice (**Figure F**). In addition, etidronate decreased mRNA abundance of *Enpp1*, *Runx2*, and *Anp* in KKAy-IR mice (**Figure F**). Similar to calcification, cardiac dysfunction was attenuated by etidronate in KKAy-IR mice, as evidenced by echocardiography

**Figure F.**
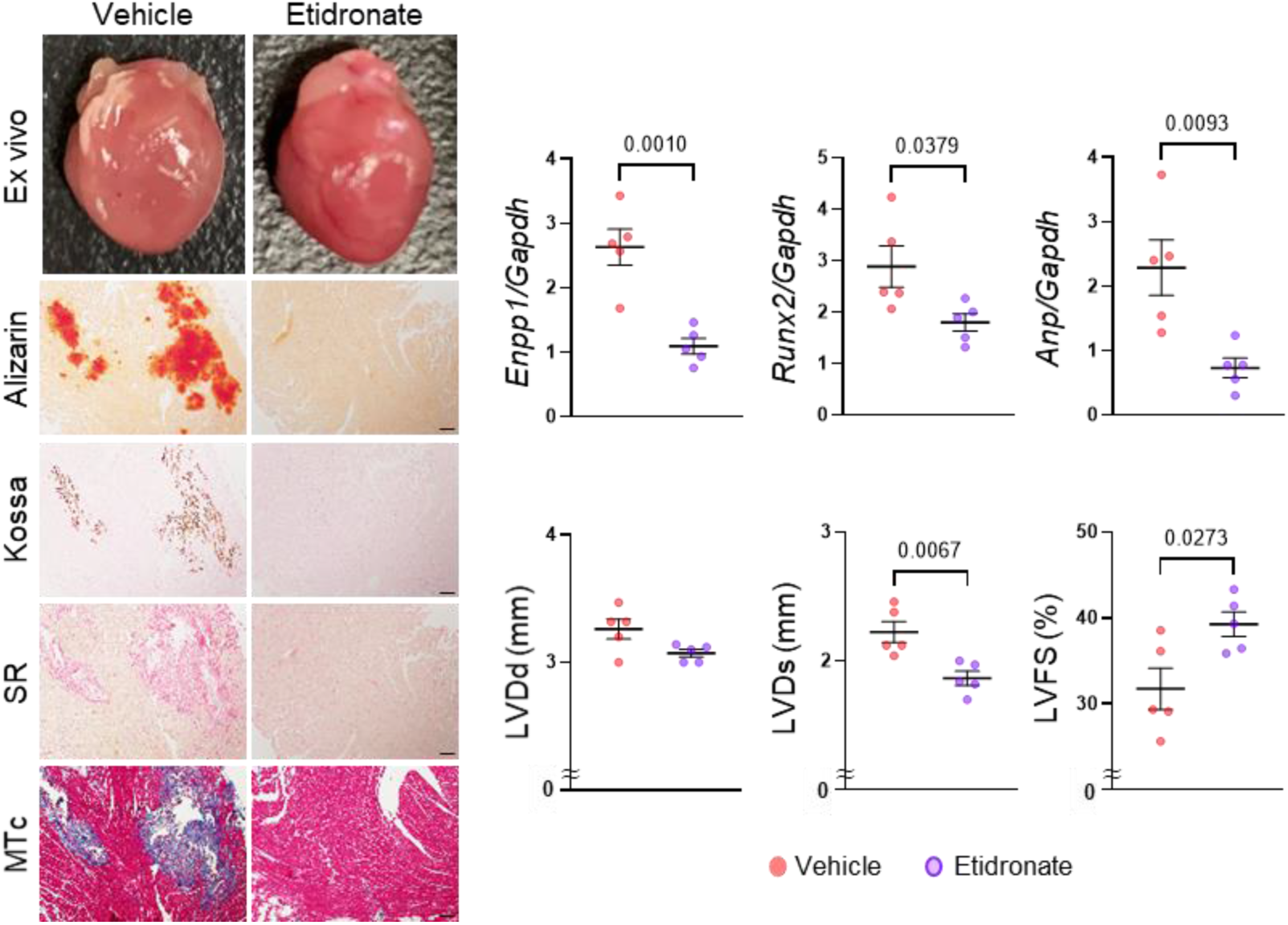
Representative gross appearance of whole hearts and heart sections stained by Alizarin red, von Kossa, SR, and MTc in vehicle or etidronate-injected KKAy mice at 14 days after IR diet (Scale bar: 100 μm). Quantitative analysis of relative *Enpp1*, *Runx2*, and *Anp* mRNA abundance, and LVDd, left ventricular end-systolic dimension (LVDs), and LVFS in vehicle or etidronate-injected KKAy mice at 14 days after IR diet. n=5 per group.

(**Figure F**). On the other hand, echocardiography demonstrated that LVDd, LVDs, and LVFS did not differ between injection in KKAy mice fed normal diet. In addition, etidronate had no significant effect on cardiac function in WT mice regardless of diets (**Figure G**).

**Figure G.**
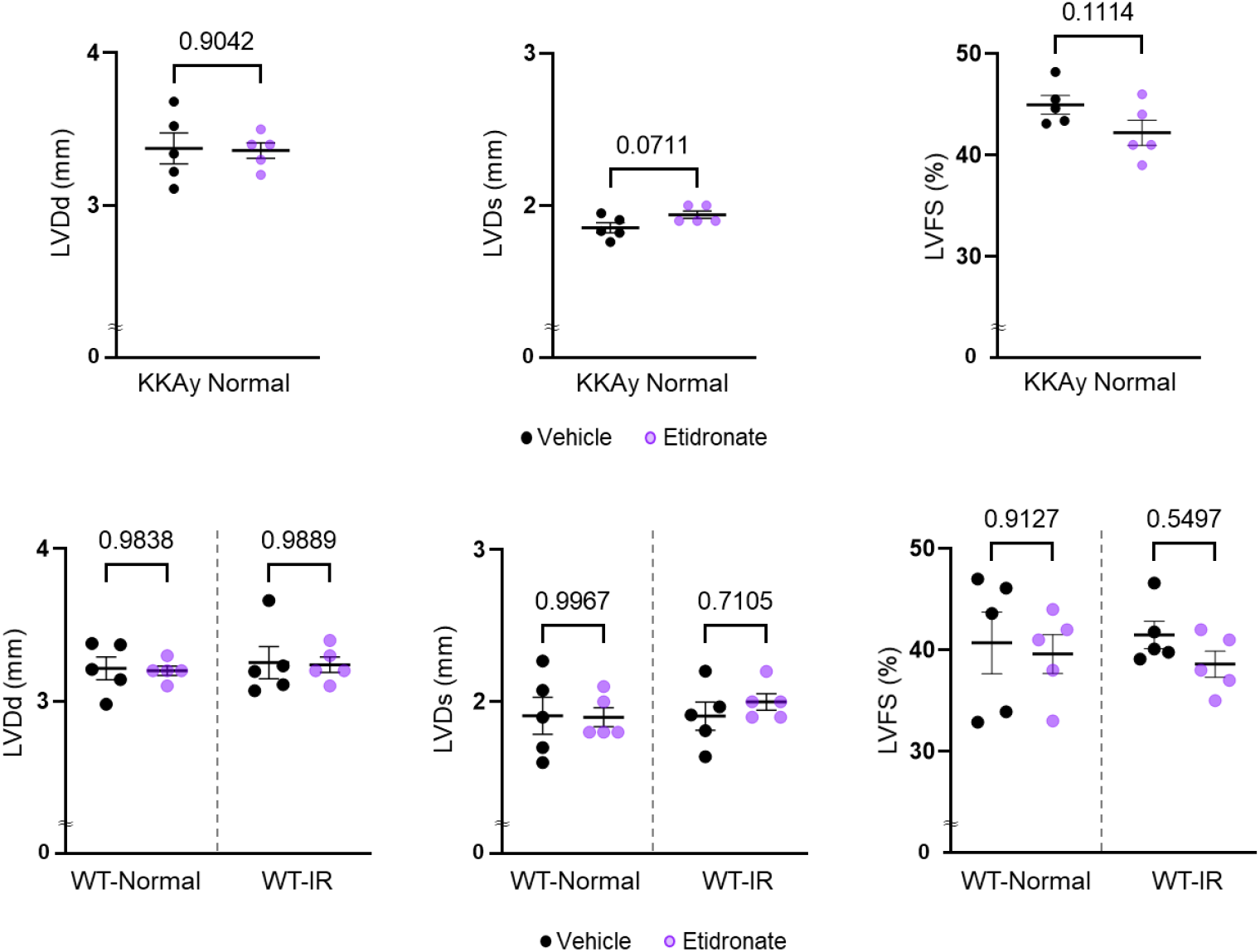
LVDd, LVDs, and LVFS in vehicle or etidronate-injected KKAy mice at 14 days after normal diet, and vehicle or etidronate-injected WT mice at 14 days after normal or IR diet. n=5 per group.

Several studies have shown that iron deficiency causes and augments heart failure through the dysregulation of ryanodine receptor 2 (*Ryr2*) and mitochondrial respiration.^3–5^ In addition, since an association of Wnt/β-catenin signaling in cardiomyopathy is reported, Wnt/β-catenin is a potential mechanism of iron deficiency induced heart failure.^6^ Then, we assessed molecules related to these mechanisms. Cardiac *Ryr2* mRNA abundance was not altered by IR diet in both genotypes (**Figure H**). Western blot analysis revealed that mitochondrial electron transfer chain complexes I-V were not different among groups. β-catenin was not changed by IR diet in both genotypes (**Figure H**). Therefore, it remains to be clarified by which iron deficiency leads to heart failure in KKAy mice.

**Figure H.**
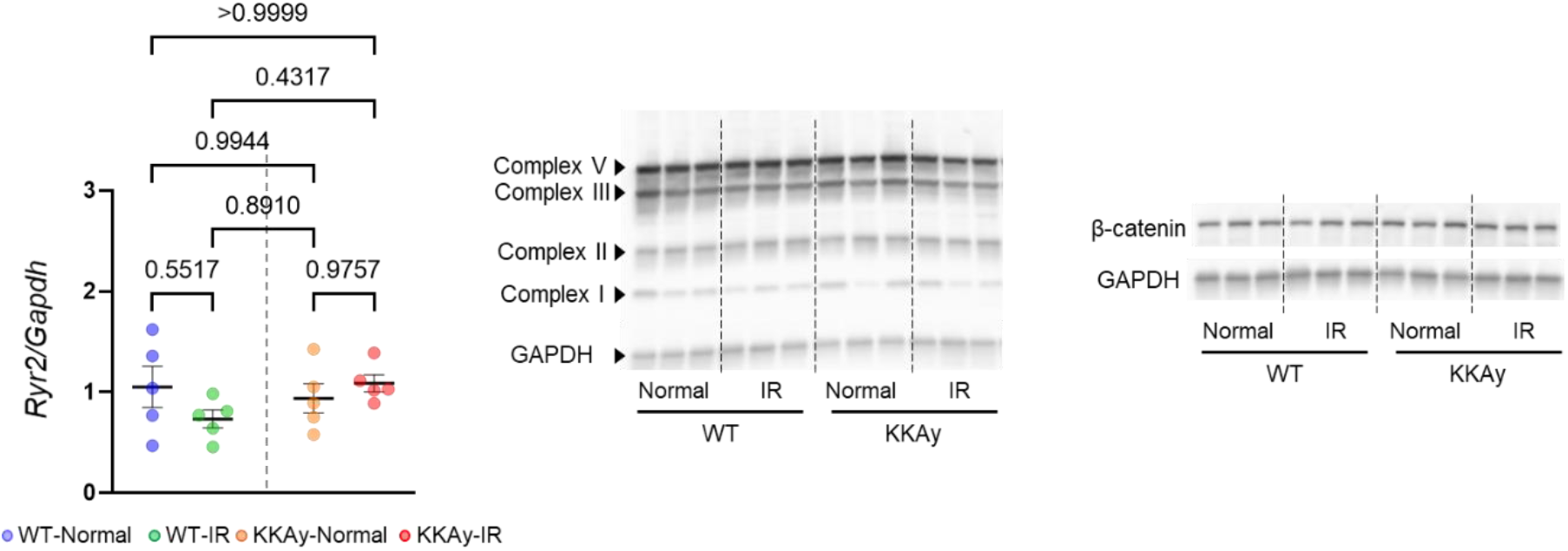
Quantitative analysis of relative ryanodine receptor 2 (*Ryr2*) mRNA abundance and Western blot analyses for **(B)** mitochondrial electron transfer chain complexes I-V, and **(C)**β-catenin in the heart of WT and KKAy mice at 12 weeks after normal or IR diet. n=5 per group.

The present study found the development of cardiac calcification and worsening cardiac function in KKAy mice by IR diet. Cardiac *Enpp1* mRNA was upregulated in KKAy-IR mice. A previous study showed that cryo-injury induced ectopic cardiac calcification with an increase of ENPP1 in mice.^7^ Thus, ENPP1 may contribute to the development of cardiac calcification. Although a number of studies have reported various molecular mechanisms by which iron deficiency leads to heart failure, it remains to be clarified how iron deficiency leads to heart failure and cardiac calcification through ENPP1 in metabolic syndrome. Importantly, our previous study reported beneficial effects of IR diet on renal damage.^8^ Therefore, the impact of IR diet may vary by diseases. It would be interesting to investigate the molecular basis of the divergent roles of iron in cardiovascular-renal diseases.

In conclusion, the present study is the first to reveal the development of heart failure along with cardiac calcification by iron deficiency in metabolic syndrome model mice. This mouse model may provide insights into understanding the complex mechanism of cardiac calcification and heart failure.

## Non-standard Abbreviations and Acronyms

Anp: atrial natriuretic peptide
BUN: blood urea nitrogen
Col1: collagen type I
Cre: creatinine
Enpp1: ectonucleotide pyrophosphatase/phosphodiesterase-1
IR: iron-restricted
Runx2: runt-related transcription factor
2 WT: wild type

## Acknowledgement

We acknowledge the technical assistance of Sachi Ito and Junko Kawata. We also thank Akari Furuse for English proofreading.

## Methods

### Mice

All experimental procedures were approved by the Animal Research Committee of Hyogo College of Medicine (protocol #16-017, #19-014). All of the animal experiments were performed in accordance with National Institutes of Health guidelines. Mice were housed in a temperature-controlled facility with a 12hr light/dark cycle and had free access to food and water.

Four-week-old male WT C57BL/6J and KKAy mice were purchased from Japan CLEA. Mice were fed either a normal diet or IR diet (#F2FeDD, Oriental Yeast Co. Ltd., Japan). Mice were subsequently terminated at 3, 7, 14 days after IR diet, and 12 weeks after normal or IR diet, and blood and hearts were harvested (n=6 per group for 3-14 days, 10 per group for 12 weeks). Etidronate (2.0mg/kg/day, #P5248, Sigma-Aldrich, St. Louis, MO, USA) was administered intraperitoneally into 4-week-old male WT and KKAy mice fed either a normal or IR diet for 14 days. Following 14 days of etidronate injection, mice were euthanized to harvest hearts (n=5 per group).

### Hematologic parameters

Blood samples were collected in the fed state. Peripheral blood cell count was measured by an automatic cell count analyzer (Pentra 60 LC-5000, Horiba, Kyoto, Japan). Blood glucose and serum creatinine concentrations were determined by enzymatic methods. Serum blood urea nitrogen concentrations were measured by the urease-glutamate dehydrogenase-UV method.

### Echocardiography

Cardiac functions, such as left ventricular end-diastolic dimension (LVDd), end-systolic dimension (LVDs), and functional shortening (LVFS), were assessed using a high frequency ultrasound system (Vevo 2100 with MS400, FUJIFILM VisualSonics, Inc., Toronto, ON, Canada). During echocardiography, a mouse was placed on a heated platform and anesthetized by isoflurane.

### Histological analysis

Heart tissues were fixed with 4% paraformaldehyde, embedded in paraffin, and cut into 4-μm-thick sections. Picrosirius red staining, von Kossa staining, Alizarin red staining, and Masson’s trichrome staining were performed using serial heart sections.

### Real-time quantitative RT-PCR

Total RNA was extracted from the heart using TRIzol reagent (Thermo Fisher Scientific, Rockford, IL, USA) as reported previously.^8^ Total RNA was reverse transcribed into cDNA using random primers (Thermo Fisher Scientific). Real-time PCR reactions were performed using a QuantStudioTM 12K Flex Real-Time PCR System with TaqMan Universal PCR Master Mix and TaqMan Gene Expression Assays (probes, Thermo Fisher Scientific). Following probes were used: *Anp* (assay ID Mm01255747_g1), *Col1* (assay ID Mm00801666_g1), *Enpp1* (assay ID Mm00501097_m1), *Runx2* (assay ID Mm00501584_m1), and glyceraldehyde-3-phosphatedehydrogenase (*Gapdh*) (assay ID Mm 99999915_g1). ΔΔCt method was used for mRNA quantification with the normalization using *Gapdh*.

### Western Blot Analysis

Cardiac proteins were extracted using freeze crusher (SK-100, Tokken Inc., Chiba, Japan) and cell lysis buffer (#9803, Cell Signaling Technology, MA, USA). Protein lysate was separated by SDS-PAGE and transferred onto polyvinylidene fluoride membranes. Protein abundance was detected by an enhanced chemiluminescence kit (#32106, Thermo Scientific, IL, USA). Anti-mitochondrial total oxidative phosphorylation antibody cocktail including antibodies against complexes I-V (#ab110413, Abcam; dilution 1:1000), anti-β-catenin (#9587, Cell Signaling Technology; dilution 1:1000), and anti-GAPDH (#2118, Cell Signaling Technology; dilution 1:1000) antibodies were used.

### Statistical analysis

All values are described as the mean±SEM or the median and 25th/75th percentiles. The assumption of normality and equal variance were assessed by Shapiro-Wilk and Brown-Forsythe tests, respectively. Student’s t- or Two-way ANOVA followed by Holm-Sidak test was applied to data confirmed the normality and homogenous variation. Kruskal-Wallis test followed by Dunn’s test was used for multiple group data not passed the normality test. Survival rate was assessed by the log-rank test. P<0.05 was considered statistically significant. Statistical analyses were performed by GraphPad Prism 9.

